# *Bacillus subtilis* spore surface display of photodecarboxylase for the transformation of lipids to hydrocarbons

**DOI:** 10.1101/2020.08.30.273821

**Authors:** Marianna Karava, Peter Gockel, Johannes Kabisch

## Abstract

The display of enzymes on the surface of spores allows the rapid and very simple biotechnological production of immobilized enzymes. Here we describe the development of a *Bacillus subtilis* spore display platform and its application to produce hydrocarbons from lipids obtained from the oleaginous yeasts *Yarrowia lipolytica, Cutaneotrichosporon oleaginosus* as well as olive oil.

Lipid hydrolysis was examined in a bienzymatic one-pot cascade using a commercially immobilized lipase (RO lipase) as well as spores with and without additional heterologous lipase expression. Decarboxylation of the released fatty acids was achieved displaying a photodecarboxylase (*Cv*FAP) on the spore surface. Differences in composition of the formed hydrocarbons were observed depending on the lipids source. Using 3D printed lighting equipment titers of up to 64.0 ± 5.6 mg/L hydrocarbons were produced.

## Introduction

The expanding global economies and population growth have increased the worldwide energy consumption. The main source of energy used to cover the rising demands is derived from fossil fuels, stadily leading to their depletion concurrently having severe environmental impacts. To address the problem of fossil fuels exhaustion and fulfill the world wide energy demands, a transition to environmentally benign and sustainable energy sources has been initiated.

For decades extensive research has been done on valorization of waste oils or vegetable oils for the production of fatty acid methyl esters and fatty acid ethyl esters (FAMEs and FAEEs) that are used as biodiesel [1–3]. Due to the technical challenges faced upon production and use of biodiesel, research has been directed toward the production of advanced biofuels from microorganisms [4]. Among a large variety of microbes, oleaginous yeasts and algae stand out due to their ability to accumulate significant quantities of lipids which could be further utilized as feedstock for the production of petroleum-like biofuels [5–8]. Several approaches have been recommended for *in vivo* production of hydrocarbons which require extensive metabolic engineering [9–11]. However very often these approaches are restricted by the low titers of the biotransformation owing to increased cellular metabolic burden and side reactions which require appropriate engineering strategies [12]. Thus it is of great interest to develop processes for the bio based production of HCs while bypassing the reconstruction of metabolic pathways.

In 2017, Beisson and coworkers identified a novel photoinducible enzyme derived from the alga *Chlorella variabilis* [13]. This flavin dependent enzyme named fatty acid decarboxylase (FAP) catalyzes the irreversible conversion of free fatty acids to the corresponding n-1 alkanes or alkenes under blue light. Since the first report several approaches have been developed employing *Cv*FAP for the production of HCs [14–16]. Hollman and coworkers suggested the reconstitution of a bienzymatic cascade consisting of a hydrolase and *Cv*FAP for the production of HCs from natural triglycerides [17]. Here we report for the first time the *in vitro* production of drop-in biofuels from natural oils using spores from *Bacillus subtilis* and employing the spore surface display technology.

The bacterium *B. subtilis* under starvation conditions undergoes a cellular differentiation process known as sporulation which is completed with the formation of a spore. Spores due to their dehydrated core are metabolically inert entities. The core is surrounded by a modified peptidoglycan layer called the cortex which in turn is encased by a multilayer proteinaceous coat [18]. The physicochemical durability of the spores together with their biological stability make them promising carriers for protein immobilization. The most common method employed for protein display on the spore surface is by fusion protein technology. The protein of interest is genetically fused to one of the anchoring proteins comprising the spore coat and therefore assembled to the spore surface. When spore formation is completed, cell lysis facilitates release of the recombinant spore to the environment. Compared to other time consuming immobilization methods where carriers are involved, spore surface display has the benefit of protein expression with simultaneous immobilization, thereby drastically reducing the cost and timescales of the biocatalyst production. Obtaining immobilized enzymes requires only cultivation followed by centrifugation. Hence, spores fulfill the economic requirements for industrial operation. This is further supported by the fact that the production of spores for spore display does not differ in principle from well-established, low-cost processes of Bacillus spores as biopesticides [19]. Next, one major advantage of spore surface display is the preservation of genotype-phenotype even under harsh conditions. Therefore spores have been exploited for the construction of protein/variant libraries [20]. Furthermore, since the intracellular formation of the spore coat is followed by the autolysis of the mother cell, the requirement for secretion and thus for optimizing signal peptides for heterologous proteins is surpassed [21]. With these advantages of de-skilled production and the potential for a drop-in scale-up in mind, this publication describes a first proof-of-principle, for renewable production of HCs from natural lipids using spore display.

## Experimental section

*Cutaneotrichosporon oleaginosus* oil was kindly provided by the group of Prof. Dr. Brück TU Munich its production is described in [22]. *Yarrowia lipolytica* oil was produced in our group (unpublished results). Olive oil originating from Greece was kindly provided by Andreas Karavas.

### Media and growth conditions

Sporulation was induced by cultivation in 2x Schaffers glucose (2xSG) medium [16 g/L Nutrient broth No. 4, 2 g/L KCl, 0.5 g/L MgSO_4_, 1 mM Ca(NO_3_)_2_, 10 μM MnCl_2_, 1 μM FeSO_4_, 0.1% D-glucose (Sigma-Aldrich, Germany)] [23]. Briefly *B. subtilis* strains were inoculated into 20 mL Lysogeny broth (LB) composed of 10 g/L tryptone, 5 g/L yeast extract, 5 g/L NaCl (LB5; Carl Roth, Germany) supplemented with 1% D-glucose, with agitation (200 rpm) at 37 °C containing the antibiotic zeocin (InvivoGen, San Diego, CA, USA) at a concentration of 20 μg/mL. Strain Bs02039 harboring high copy plasmid of the *Cv*FAP fusion protein was cultivated in media containing kanamycin (Carl Roth, Germany) at final concentration of 50 μg/mL. Cell cultures were grown overnight at 30 °C with agitation (200 rpm). Cells from the overnight culture were harvested by centrifugation and inoculated into 2 L Erlenmeyer flasks containing 400 mL prewarmed 2xSG medium. Cells were grown at 30°C with agitation (200 rpm) for 48 h.

### Spores purification

For spore purification, after 48 h of cultivation the 400 mL cultures were harvested by centrifugation at 3,400 x g for 15 min. The spores were treated with lysozyme for removal of residual cells. Briefly, spore pellet was resuspended in 80 mL of freshly prepared lysozyme (75 μg/mL) and incubated at 30 °C with agitation (200 rpm) for 45 min. The lysed cells were centrifuged at 3,488 x g for 15 min at 4 °C. Pellet was resuspended in 80 mL of Tris-Buffer (50mM, pH 8) and centrifuged once again. This washing step was repeated one more time. Afterwards, spore pellet was resuspended in 4 mL Tris-Buffer (100 mM, pH 8.5 unless noted otherwise) and OD_600 nm_ was measured. For the spore dry weight (SDW), 500 μL-1 mL of the spore stock was transferred to a microcentrifuge tube with defined weight. The suspension was centrifuged at 16,000 x g for 10 min. Supernatant was removed while the spore pellet was placed in a dessicator filled with silica beads for 48 h. Afterwards, SDW was determined gravimetrically.

### Plasmid construction and *B. subtilis* transformation

For display of *Cv*FAP on spores, the coding sequence of the gene was synthesized codon optimized for *B. subtilis* and cloned in pET28 vector. Sequentially the gene fragment was introduced in the appropriate vector by employing a modified SLiCE method as described by [24]. The coding gene was fused in frame to the C-terminal coding end of the gene *cotY* downstream of a flexible linker (2 repeats of Gly-Gly-Gly-Gly-Gly-Ser). C-terminally the protein harbored a 6xHis tag. The fusion was expressed under the control of sporulation inducible promoter P*cotYZ*.

The resulting plasmids were isolated from *E. coli*, verified via sequencing and transformed into *B. subtilis* via natural competence as described previously [32]. For strain Bs02035, the construct was genomically integrated into the *pksX* locus via homologous recombination. Strain Bs02039 contains a single copy of the fusion protein and the promoter integrated in the *pksX* locus and additionally a high copy plasmid with the same construct. A list of all plasmids used in the present work can be found in Table 1.

**Table 1:** List of strains employed in the present study.

| Strain | Genotype | Resistance | Reference |
| --- | --- | --- | --- |
| *Bacillus subtilis* KO7 | $\Delta nprE \Delta aprE \Delta epr$<br>$\Delta mpr \Delta nprB \Delta vpr$<br>$\Delta bpr$ | no | *Bacillus Genetic*<br>*Stock Center ID*<br>1A1133 |
| Bs02003 | $\Delta sacA::(Zeo^R,$<br>$P_{xyIA}-cre, P_{spac}-comS,$<br>$lacI), \Delta cwID::lox72,$<br>$\Delta sleB::lox72$ | zeo<sup>R</sup> | Previous study [40] |
| Bs02035 | $\Delta sacA::(Zeo^R,$<br>$P_{xyIA}-cre, P_{spac}-comS,$<br>$lacI), \Delta cwID::lox72,$<br>$\Delta sleB::lox72$<br>$\Delta pksX::(P_{cotYZ}-cotY-li$<br>$nker-CvFAP)$ | zeo<sup>R</sup> | This work |
| Bs02039 | Bs02035 +<br>replicative plasmid<br>p02030 | Kan<sup>R</sup> | This work |
| Bs95002 | Bs02003 +<br>replicative plasmid<br>p95002 | Kan<sup>R</sup> | BSc thesis of Peter<br>Gockel |

### Decarboxylation of palmitic acid

All reactions were performed in 5 mL transparent glass vials (ND13, Sarstedt). Decarboxylation reactions were performed in biphasic systems containing n-hexane as organic solvent. Standard reaction volume was 600 μL containing spores of OD_600 nm_~40 (4 mg/mL of SDW) resuspended in 300 μL Tris-HCl buffer (100 mM, pH 8.5) and 300 μL n-hexane containing 15 mM palmitic acid and 1 mM of dodecane. The mixture was vortexed for 5 sec in order to form stable emulsions. The vials were placed in a rotor covered with a custom made drum with blue LEDs (S1 Figure and CAD-files available at https://gitlab.com/kabischlab.de/led-labware-cvfap). Vials were rotated at 70 rpm and illuminated continuously with blue light (130 mAmp, 4 V, 68 μmol quanta m^-2^ s^-1^) for 20 h at 30 °C unless noted otherwise. Reactions were terminated by switching off the light source and extraction of the HCs. As a control for detection of native fatty acid decarboxylase activity, equal amounts of spores from an untransformed strain (Bs02003) were employed.

### Biotransformation of Triglycerides to HCs in one-pot reaction

One-step biotransformation (1 mL): The final reaction consisted of 10 μL triolein from stock solution with concentration of 0.2 M in n-hexane or 25 μL oil (*C. oleaginosus, Y. lipolytica*, olive oil), 50 mg of immobilized lipase from *Rhizopus oryzae* (≥300 U/g, Sigma-Aldrich) and 990 or 975 μL of spore suspension in 100 mM Tris-HCl, pH 8.5. The spore suspension in the reaction had OD_600 nm_~120 corresponding to 16 mg/mL SDW. Reactions were placed in the rotor and were incubated at 30 °C with constant stirring at 70 rpm for 24 hours.

Two-step biotransformation (1 mL): The first step of the reaction consisted of 10 μL triolein from stock solution with concentration of 0.2 M in n-hexane, 50 mg of immobilized lipase from *Rhizopus oryzae* (≥300 U/g, Sigma-Aldrich) or alternatively spores from strain Bs95002 displaying the lipase from *Rhizopus oryzae* with OD_600 nm_~ 160 (21 mg/mL SDW). The reaction volume in the first step was 700 μL. The hydrolysis reaction took place at 30 °C with agitation at 70 rpm for 48 hours. Afterwards, 300 μL of Bs02039 spore suspension with final OD_600 nm_~ 120 (16 mg/mL SDW) were added in the mixture and the reaction was further incubated for 24 hours in the presence of blue light.

In both types of biotransformation, the reaction vials were vortexed for 5 sec prior to incubation. Reactions with *Cv*FAP were started by activating the LED light source (130 mAmp, 4 V, 68 μmol quanta m^-2^ s^-1^). Reactions were terminated by removing the samples from the light source and storing them in −20 °C. Following, extraction of HCs and GC analysis were performed. As negative control, strain Bs02003 was employed. To elucidate the putative hydrolytic activity of spores, reactions without the addition of lipase were performed. All reactions were carried out in triplicates. Data analysis was performed with R version 3.6.3 (2020-02-29).

### Extraction of HCs

For the extraction of HCs, reactions were transferred to 2 mL extraction tubes containing an amount of glass beads (0,25-05 mm). When needed, reactions were supplemented with n-hexane so the minimum volume of the organic phase was between 150 and 300 μL. Subsequently, the extraction tubes were vortexed vigorously using a ball mill (Mixer Mill MM 400) at 30 Hz for 15 min. After the extraction, mixtures were centrifuged at 16,000 x g for 15 min and the upper organic phase was transferred to a glass vial for GC analysis.

### Analysis of HCs with gas chromatography (GC)

Detection of HCs was performed by gas chromatography as described by [14]. The samples were analyzed with a Shimadzu Nexis GC 2030, on a Shimadzu SH-Rxi-5MS column (30 m, 0.25 mm, 0.25 μm) and detected by FID. The temperature of inlet and FID were set to 250 °C and 310 °C, respectively. The linear velocity of hydrogen was set to 50 cm/s. Split was set to 1. Column oven temperature program: Temp. 90 °C, hold 5 min; Rate 15, final temp. 190 °C; Rate 2.0, final temp. 200 °C, hold 1 min; Rate 0.5, final temp. 202.5 °C, hold 1 min; Rate 20, final temp. 300 °C, hold 5 min.The analytical GC grade standards undecane, tridecane, pentadecane, heptadecane and the C8-C20 alkane standard solution were purchased at Sigma-Aldrich. Quantification of 7-pentadecene, 8-heptadecene and 6,9-heptadecadiene were performed according to [25]. Data processing was done with LabSolutions 5.92 and R version 3.6.3 (2020-02-29).

## Results & Discussion

### Strain engineering for improved enzymatic activity

Among the different proteins that form the spore coat, in the current work CotY was selected which is located on the most outer layer of the spore coat, called the crust. The selection of this protein was based on its location on the spore coat and its abundance according to previous findings [26]. The gene encoding *Cv*FAP was fused in frame to the C-terminal coding end of the gene *cotY*. The two genes were bridged by a glycerin linker of 12 amino acids. The fusion was constructed this way by taking into account the protein structure of *Cv*FAP, which revealed that the N-terminal site of the protein forms a long flexible helix (PDB:5NCC) [13]. At the C-terminus *Cv*FAP was fused to a 6xHis-tag to allow for its localization on the spore coat. The expression of the fusion gene was regulated by the native promoter of *cotY*which is naturally induced during sporulation. To avoid germination and subsequent outgrowth of the spores, all strains employed harbor deletion of *cwID* [27] and *sleB* [28] genes.

Toward the production of more efficient spore based catalysts, we anticipated that an increase in the gene copy number of the fusion would lead to higher enzyme concentration and activity. Therefore a high copy plasmid [29] was constructed expressing the *Cv*FAP fusion gene. The effectiveness of the high copy strain Bs02039 was compared to strain Bs02035 which harbors a single copy of the same construct genomically integrated. Both strains were compared to the untransformed strain Bs02003. The constructed strains were tested for their efficiency to convert palmitic acid to pentadecane in a biphasic system containing n-hexane.

As expected the highest enzymatic activity was obtained for strain Bs02039 which harbors a high copy plasmid, whereas only trace amounts of pentadecane were detected for strain Bs02035 (Figure 1). This 19-fold increase in activity can be associated with higher protein load on the spore surface stemming from the multiple gene copies. Interestingly the overall reaction conversion did not exceed 1%. This significantly low yield is likely due to photoinactivation of the displayed enzyme. Previous studies suggested that exposure of *Cv*FAP to blue light is directly linked to the formation of radical species, leading to irreversible inactivation of the enzyme [30]. Based on the better performance of Bs02039 all further experiments were conducted with this strain.

**Figure 1:**
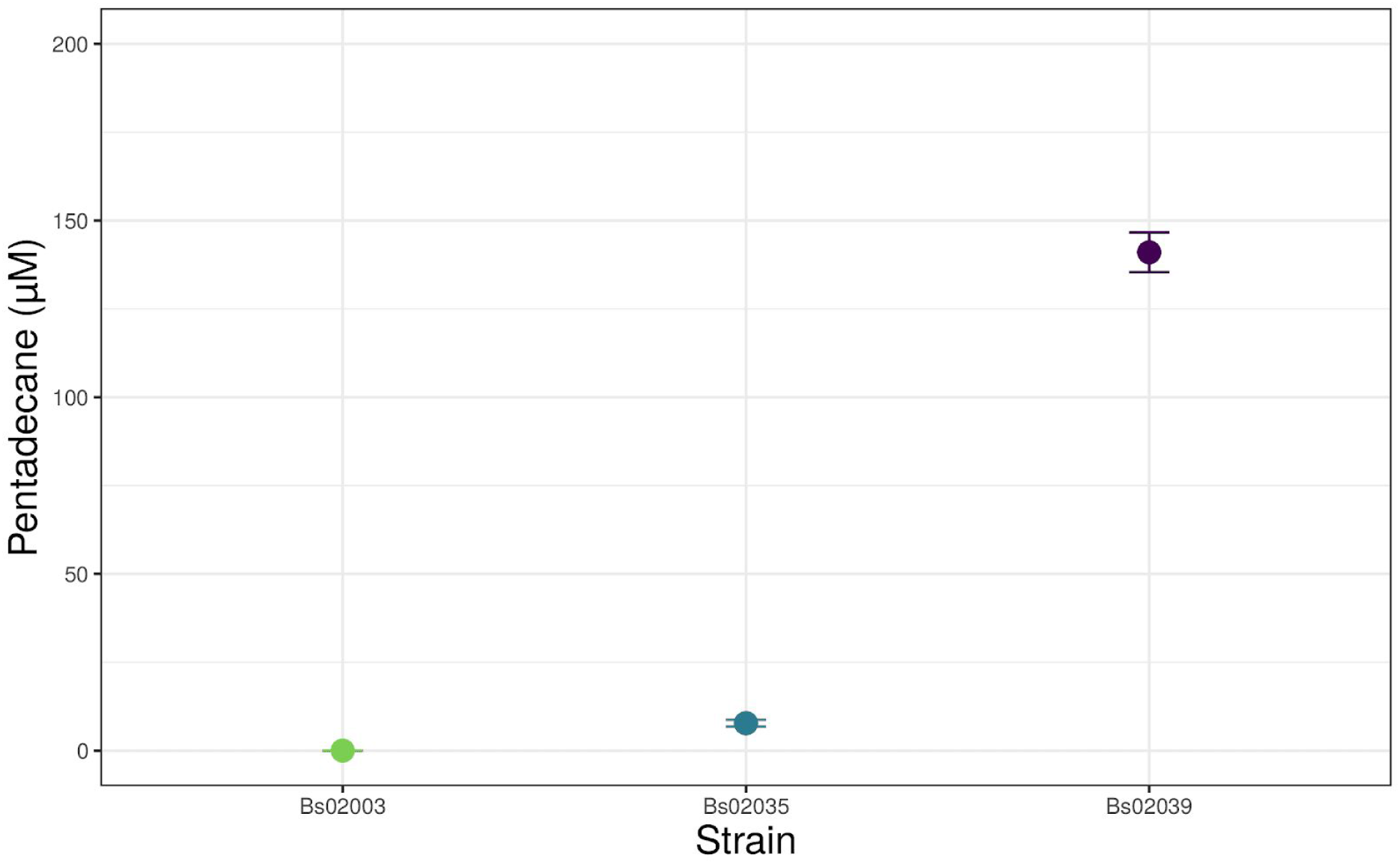
Gene dosage effect in the activity of spore displayed *Cv*FAP. Three different strains were tested: Bs02003 is the negative control, Bs02035 has a single copy of the gene fusion *cotY*-*Cv*FAP and Bs02039 has a high copy plasmid. Reaction conditions: 15 mM palmitic acid in n-hexane (50 % v/v), 3 mg of SDW, 30 °C, 20 h and blue light illumination (130 mAmp, 4 V, 68 μmol quanta m^-2^ s^-1^). Data are mean values of quadruplicates. Error bars indicate the standard deviation (n=4).

### Stability of spore catalyst under different reaction conditions

To gain deeper insight into the properties of the spore displayed *Cv*FAP, different factors with a putative effect in the catalytic activity toward decarboxylation of palmitic acid were tested.

As shown in Figure 2A, an increase in reaction temperature from 30 to 37 °C was followed by a reduction in the catalytic activity of the enzyme. An explanation for this could lie in the origin of *Cv*FAP from microalga which have optimal growth temperatures between 15 °C and 30 °C [31]. The pH profile of the spore displayed *Cv*FAP (Figure 1B), results in a maximum product formation at pH 9.0 while almost 60% decrease in the pentadecane titers were observed at pH 7.0 and 10 respectively. Compared to the published data for the free enzyme it seems that immobilization of the catalyst slightly increased the pH optimum from 8.5 to 9.0 [13]. This change could be linked to the complex nature of the spore coat which contains different molecules including proteins and polysaccharides [32, 33]. To investigate the organic solvent-tolerance of the catalyst, three different concentrations of n-hexane were employed. Figure 1C shows that product formation decreases with the concentration of the organic phase indicating a lower stability of the catalyst in higher solvent concentrations. Next, the influence of catalyst concentration was examined. As expected, higher spore concentrations led accordingly to better reaction yield (Figure 1D). This is especially important in light of the potentially very easy scaling of spore production due to spores already being produced in bulk as biopesticides [19].

**Figure 2:**
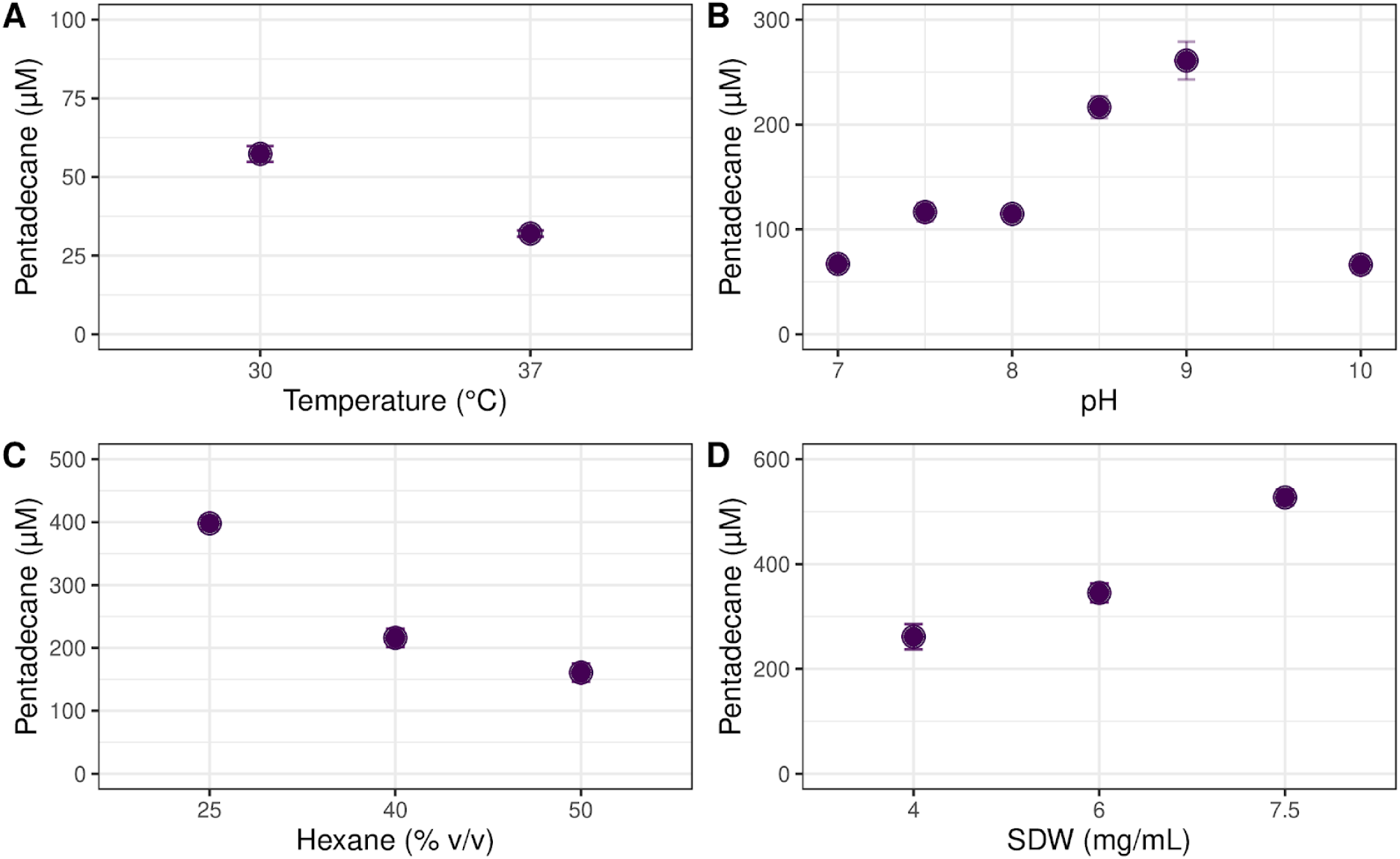
Enzymatic activity of the displayed *Cv*FAP (Bs02039) as a function of different reaction conditions. All reactions were performed using 15 mM palmitic acid in n-hexane as substrate and blue light illumination. Incubation time was 20 h. A: Reaction volume was 600 μL, catalyst (SDW) concentration was 4 mg/mL in Tris-HCl buffer (100 mM, pH 8.5), 50 % n-hexane. B: Reaction volume was 600 μL, catalyst (SDW) concentration was 4 mg/mL in Tris-HCl buffer (100 mM), 50 % n-hexane. C: Catalyst (SDW) concentration was 4 mg/mL in 300 μL Tris-HCl buffer (100 mM, pH 9.0). D: Reaction volume was 400 μL, catalyst (SDW) concentration was 4-7.5 mg/mL in Tris-HCl buffer (100 mM, pH 9.0), 25 % n-hexane. Data are mean values of quadruplicates Error bars indicate the standard deviation (n=4).

Despite our efforts to optimize the reaction conditions for *Cv*FAP, the obtained pentadecane concentrations roughly exceed 0.5 mM corresponding to ~3% conversion. By monitoring the product formation over time (S2 Figure), we observed that the reaction was terminated within 3-4 hours, thus confirming previous findings reporting on the unstable nature of *Cv*FAP [30].

### Spore mediated biotransformation of triolein to HCs by employing a bienzymatic cascade

For the production of HCs from triglycerides(TAGs), as a proof of concept a one-pot bienzymatic catalysis was performed employing triolein as substrate. The enzymatic cascade comprises a lipase and the light inducible decarboxylase CvFAP [17]. For the hydrolysis step two different variants of immobilized lipase from *Rhizopus oryzae* were tested. The one variant included the lipase displayed as a fusion protein on the surface of the spores (Bs95002) while the other variant was commercially available immobilized lipase (RO lipase). Furthermore the putative hydrolytic activity of the spores was assessed by employing spores from strain Bs02003 without heterologous expression of lipase. The decarboxylation step was carried out by spores from strain Bs02039. To investigate the optimal conditions for the biotransformation, the cascade was carried out in one-step and two sequential steps respectively.

**Figure 3:**
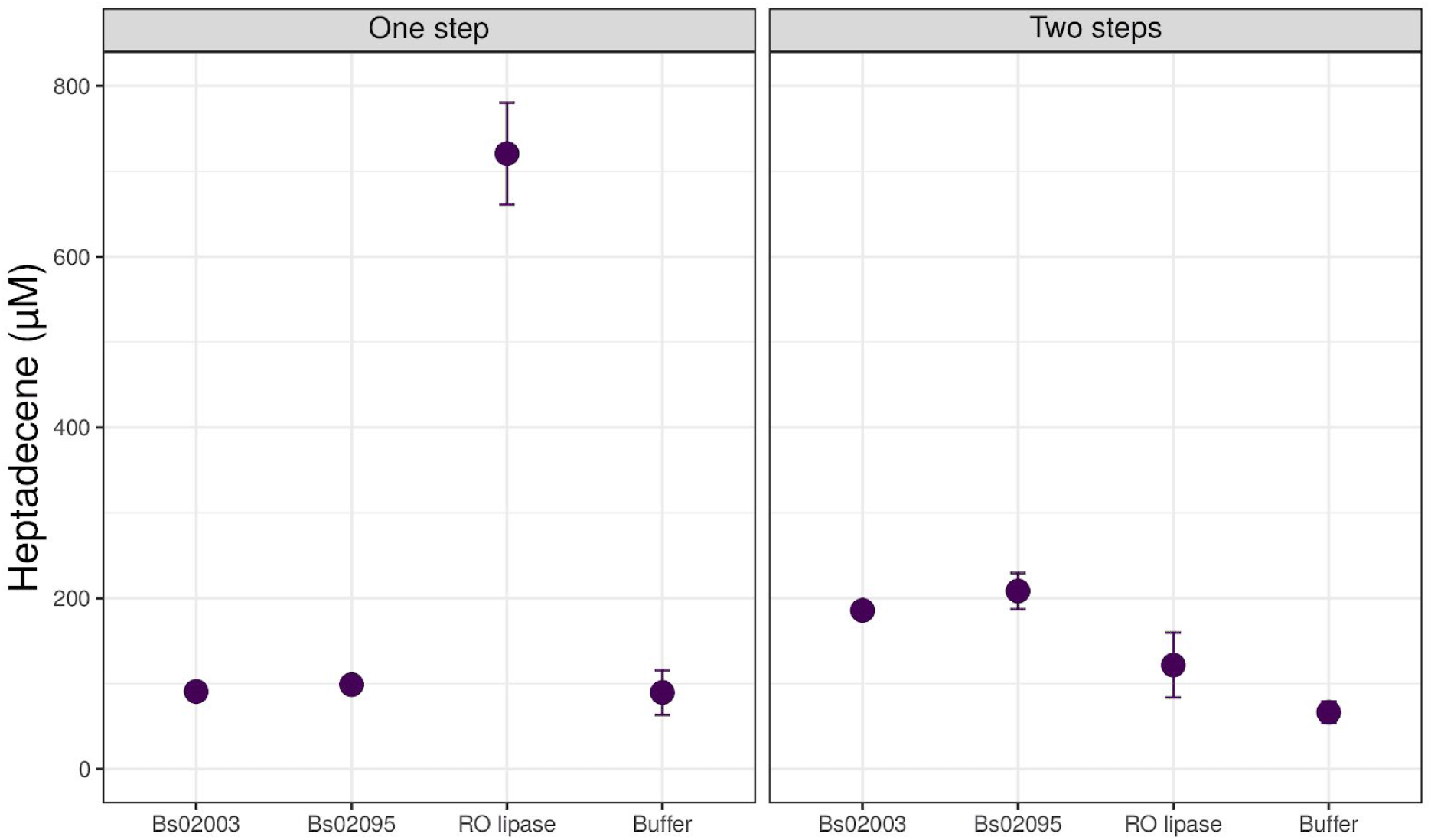
One pot enzymatic conversion of triolein to heptadecene (C17:1). The biotransformation was carried out in one and two step reactions respectively testing different lipases. One step reaction was carried out within 24 h. For the two step sequential catalytic reaction, Bs02039 spores were added after 48 h and the reaction was further carried out for 24 h. All reactions were performed in Tris-HCl buffer (100 mM, pH 8.5) using triolein as substrate and blue light illumination (130 mAmp, 4 V, 68 μmol quanta m^-2^ s^-1^). SDW concentration was 16 mg/mL. For the reactions where Ro lipase was tested, 50 mg/mL of RO lipase were added. As control for the hydrolysis reaction, spores from strain Bs02003 were employed as well as reactions without addition of lipase (WO) were prepared. Data are mean values of triplicates. Error bars indicating standard deviation (n=3).

The maximum 8-heptadecene (C17:1) formation was obtained when the hydrolysis step was catalyzed by RO lipase in one-step biocatalysis, yielding a product concentration of approximately 0.7 mM which corresponds to roughly 3.5 % conversion. The titer levels were 3-fold higher than those attained with the two-step biotransformation indicating a better synergy between the displayed *Cv*FAP and the lipase in the one-step catalysis. Upon visual inspection the addition of spores in the reaction assisted in better emulsification of the individual components, thus featuring a more uniform distribution of substrates and catalysts. The emulsifying effect of spores and cells in biphasic systems has been investigated in previous studies [34, 35].

The activity obtained from the spore displayed lipase (Bs95002) in the one-step reaction was in a similar range as for the untransformed strain Bs02003 and the control reaction, suggesting that the spore displayed lipase was not active under these conditions. Notably, the 8-heptadecene (C17:1) levels obtained for both strains in the two-step reaction were elevated compared to the one-step catalysis, suggesting that the spores retain a certain hydrolytic activity. The background hydrolytic activity presumably arises from the numerous enzymes comprising the spore coat. The spore protein LipC for example has been annotated as a lipase. However to date only a few studies regarding the substrate specificity and the role of this enzyme are available [36, 37].

### Spore mediated production of drop-in biofuels from natural oils employing *Cv*FAP

Microbial oils extracted from the oleaginous yeasts *C. oleaginosus* (Co. oil) and *Y. lipolytica* (Yali. oil) as well as olive oil were used as substrates for the production of HCs. Inspired by the results for the triolein conversion, the biotransformation was carried out in one-pot one-step bienzymatic cascade consisting of RO lipase and the spore displayed *Cv*FAP (Bs02039). As control, the reactions were performed without the addition of lipase. The produced titters of HCs are shown in Figure 4 whereas the total amount of HCs obtained for each substrate is shown in Table 2.

**Figure 4:**
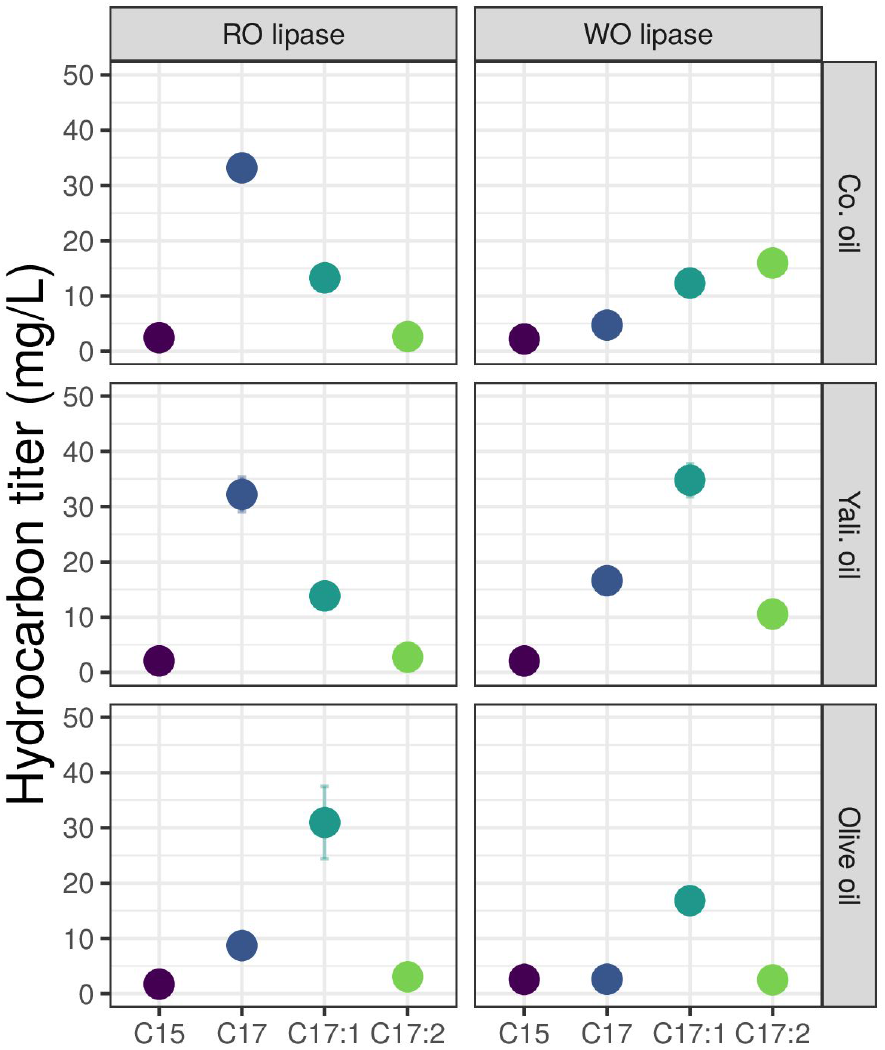
Biotransformation of natural oils (Co. oil, Yali. oil, olive oil) to HCs catalyzed by the spore displayed *Cv* FAP (Bs02039). Profile of hydrocarbons produced with (RO) and without (WO) the addition of lipase. All reactions were performed in Tris-HCl buffer (100 mM, pH 8.5) using 25 μL of oil as substrate (2.5% v/v) and blue light illumination (130 mAmp, 4 V, 68 μmol quanta m^-2^ s^-1^). SDW concentration was 16 mg/mL. For the reactions where lipase was tested, 50 mg/mL of RO lipase were added. Reaction time was 24 h. Data are mean values of triplicates. Error bars indicate the standard deviation (n=3).

**Table 2:** Titers of HCs produced from different oils with (RO) and without (WO) lipase.

| Oil | Lipase | Titters (mg/L) |
| --- | --- | --- |
| Co. oil | RO | 51.5 ± 3.6 |
|  | WO | 35.2 ± 5.2 |
| Yali. oil | RO | 50.8 ± 4.5 |
|  | WO | 64.0 ± 5.6 |
| Olive oil | RO | 44.4 ± 8.9 |
|  | WO | 24.5 ± 1.2 |

The selected oils have a fatty acid composition mainly made up of long-chain fatty acids [14, 22, 38, 39]. As shown in Figure 4, in the reactions with microbial oils as lipid substrate (Co. oil, Yali. oil) the addition of RO lipase increased production of heptadecane (C17), confirming the preference of *Cv*FAP for saturated fatty acids [17]. In contrast, without the addition of lipase (WO lipase) the hydrocarbon profile was changed in favor of unsaturated HCs, with 8-heptadecene (C17:1) and 6,9-heptadecadiene (C17:2) accounting in sum for more than 80 % and 70 % of the overall hydrocarbon fraction in each oil (S3 Figure). This change was not observed when olive oil was used as substrate, with 8-heptadecene (C17:1) being the predominant hydrocarbon produced in both reactions. The sum of produced HCs (Table 2), indicates that in the cases of Co. oil and Olive oil, the presence of RO lipase resulted in only a minor increase of the overall hydrocarbon titters while in the case of Yali. oil the highest titers were obtained in the control reaction (64.0 ± 5.6 mg/L).

No HCs were detected for any of the substrates when spores from strain Bs02003 were utilized as a control for native fatty acid decarboxylase activity (S4 Figure).

The above results suggest that the utilized oils already contained free fatty acids, potentially as the product of hydrolytic activity from native lipases or by fatty acids released during the extraction process. Since only a small shift in the overall titers was obtained with addition of lipase in the reaction, we hypothesize that the rate limiting factor in the reaction is not the substrate availability but the stability and substrate specificity of *C*vFAP.

## Conclusion

In the present work we have demonstrated a very simple, spore display based bioproduction of hydrocarbons from lipids. Catalytic activity of the photodecarboxylase *Cv*FAP was greatly increased by manipulating the gene copy number. We could show the ability of the spore displayed enzyme to produce hydrocarbons in a one-pot bienzymatic cascade. Although the produced titers are low compared to previous *in vitro* approaches, this first step has demonstrated the simplicity of producing and purifying immobilized enzymes using this technology. We are certain that our findings, future engineering of the *Cv*FAP for increased stability and process optimizations, along with the scalability of spore production will permit to improve overall yields toward a feasible application for the renewable production of hydrocarbons.

## Abbreviations

SDW: spore dry weight
TAGs: triacylglycerides
HC: hydrocarbon
Co. oil: natural oil produced by Cutaneotrichosporon oleaginosus
Yali. oil: natural oil produced by Yarrowia lipolytica
RO: *Rhizopus oryzae*
WO: without
SG: Schaffers glucose

## Acknowledgements

We thank Eva Moldenhauer for providing the *Yarrowia lipolytica* oil. MK is supported by the BMBF grant Cascade Kit (FKZ 031B0579A).

## Supplemental

**S1 Table.** Plasmids employed in the present study.

| Plasmid | Backbone | Insert | Integration site | Resistance | Reference |
| --- | --- | --- | --- | --- | --- |
| p02002 | pJET | *cwID* |  | spec | This study |
| p02005 | pJET | *sleB* |  | spec | This study |
| p02030 | pMSE4 | ORI: colE1 (Ec), repE (Bs); KanR; Resolvase beta; PcotYZ, cotY, CvFAP | - | kan | This study |
| p95002 | pMSE4 | ORI: colE1 (Ec), repE (Bs); KanR; Resolvase beta; PcotYZ, cotY, ROL(*R. oryzae* lipase) | - | kan | BSc thesis from Peter Gockel |

**Figure S1:**
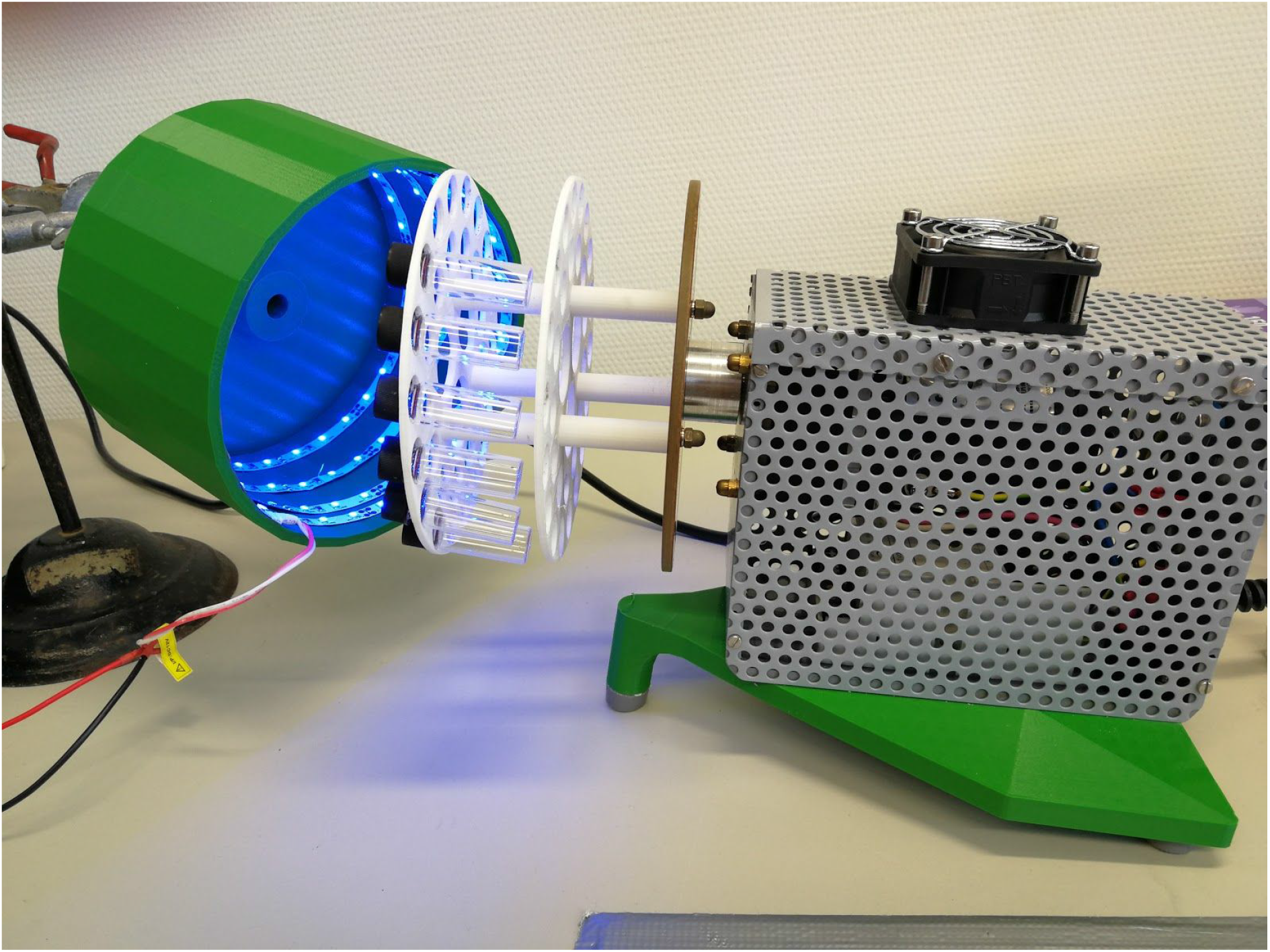
Custom made photocatalytic set up for small scale production of hydrocarbons. CAD files available at https://gitlab.com/kabischlab.de/led-labware-cvfap

**Figure S2:**
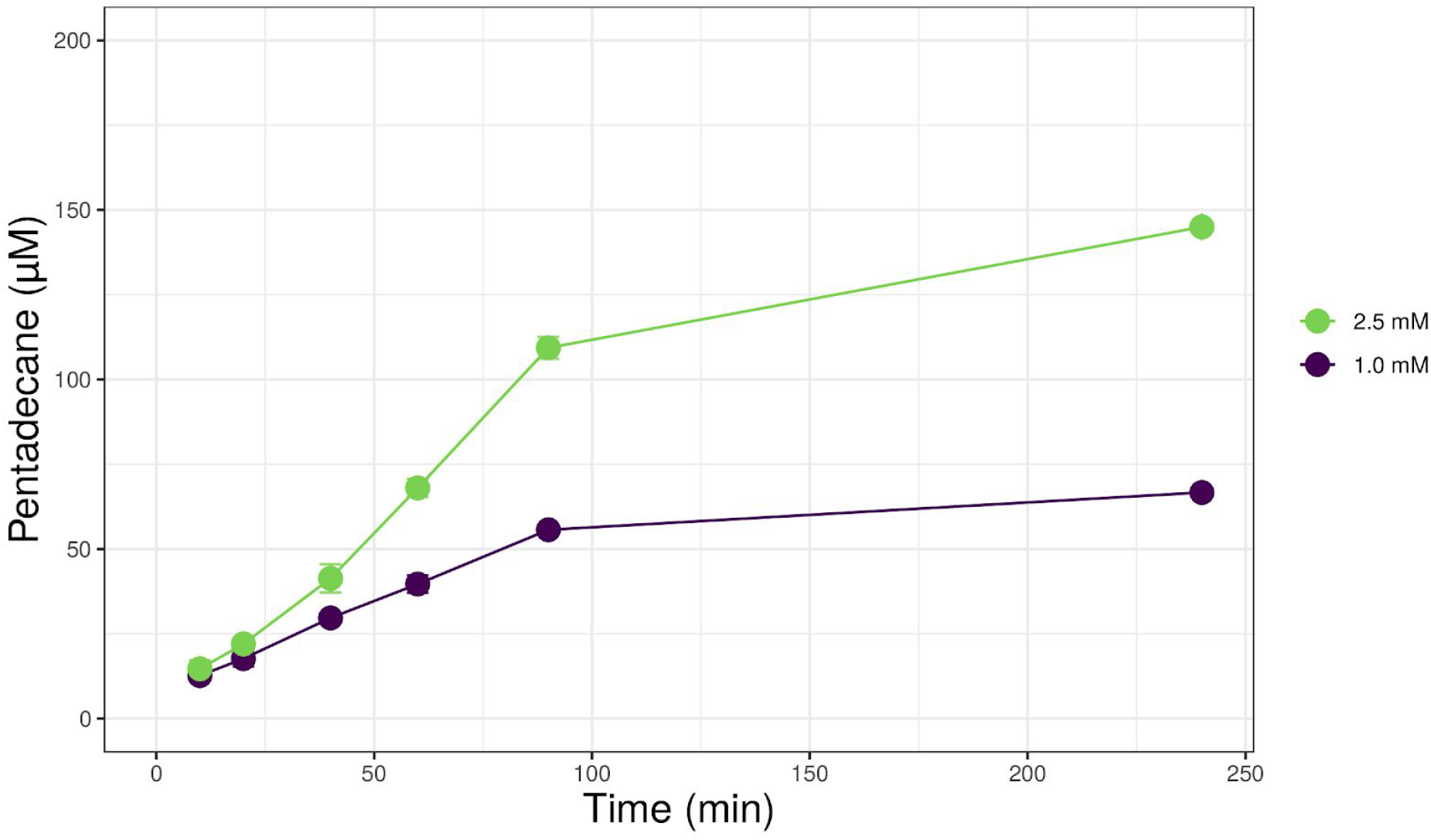
Time course production of pentadecane from *Cv*FAP using two different substrate concentration. Reaction conditions: 1.0-2.5 mM palmitic acid, 16 mg/mL of SDW, 30 °C, 70 rpm, blue light illumination (130 mAmp, 4 V, 68 μmol quanta m^-2^ s^-1^)

**Figure S3:**
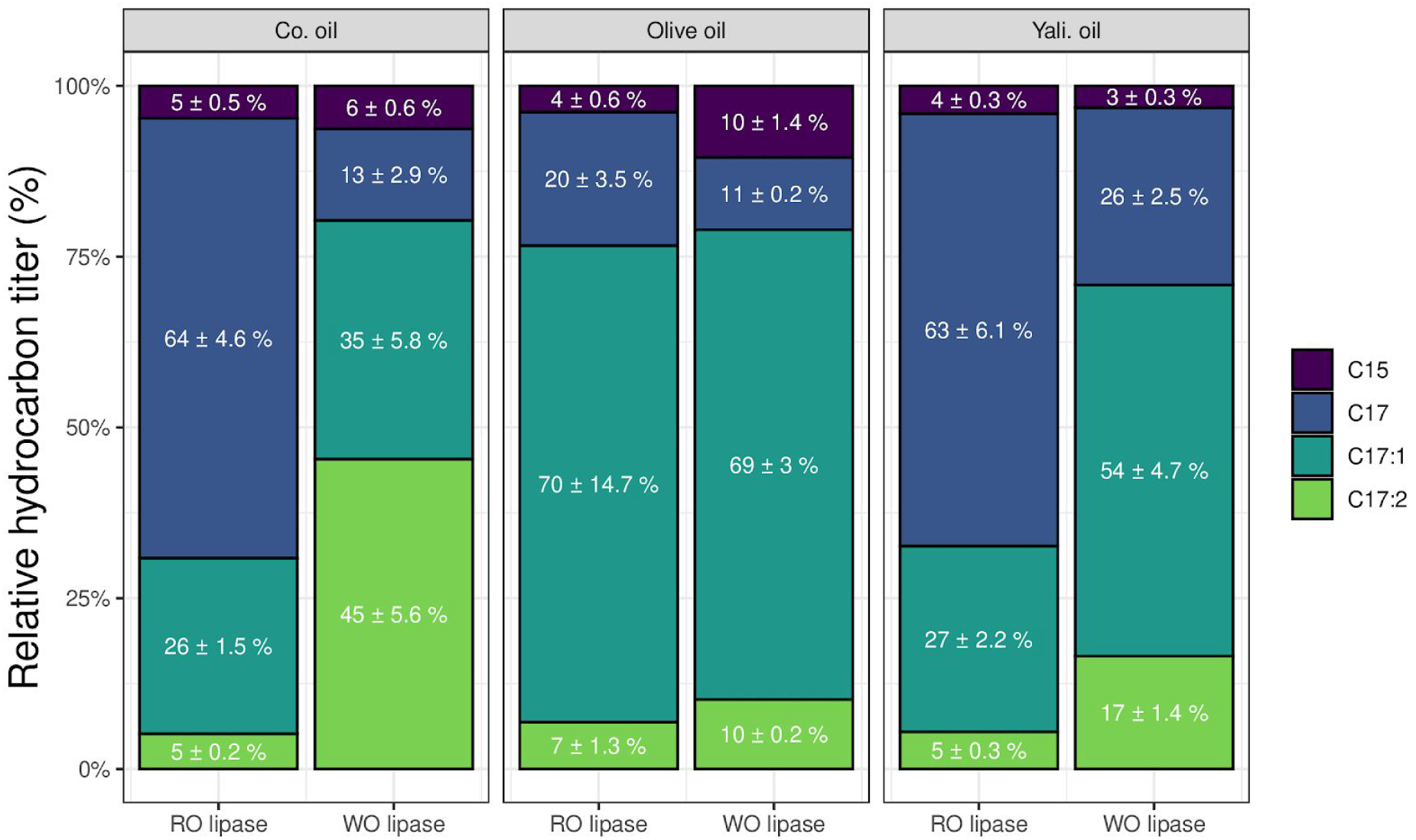
Relative composition of hydrocarbon titers in percent (%) as produced from the reaction of *Cv*FAP with different oil sources.

**Figure S4:**
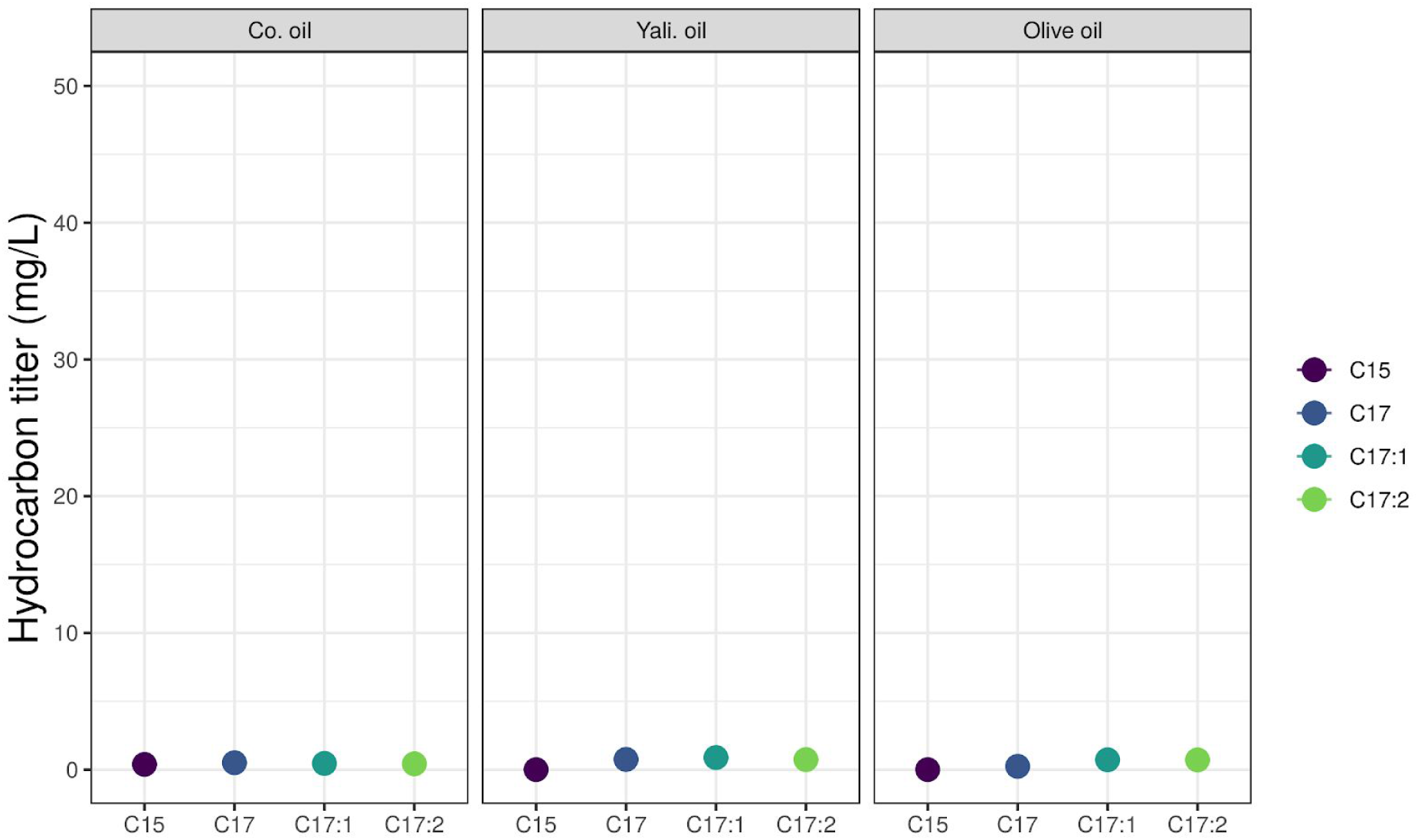
Produced hydrocarbon titers from the reaction of "wild type” BsO2OO3 spores not displaying any heterologous enzyme with different oils. Reaction conditions: 1.0-2.5 mM palmitic acid, 16 mg/mL of SDW, 30 °C, 70 rpm, blue light illumination (130 mAmp, 4 V, 68 μmol quanta m^-2^ s^-1^)

